# GSuite HyperBrowser: integrative analysis of dataset collections across the genome and epigenome

**DOI:** 10.1101/067561

**Authors:** Boris Simovski, Daniel Vodak, Sveinung Gundersen, Diana Domanska, Abdulrahman Azab, Lars Holden, Marit Holden, Ivar Grytten, Knut Rand, Finn Drabløs, Morten Johansen, Antonio Mora, Christin Lund-Andersen, Bastian Fromm, Ragnhild Eskeland, Odd Stokke Gabrielsen, Sigve Nakken, Mads Bengtsen, Alexander Johan Nederbragt, Hildur Sif Thorarensen, Johannes Andreas Akse, Ingrid Glad, Eivind Hovig, Geir Kjetil Sandve

**Affiliations:** Department of Informatics, University of Oslo, Oslo, Norway; Department of Mathematics, University of Oslo, Oslo, Norway; Statistics For Innovation, Norwegian Computing Center, Oslo, Norway; Institute for Medical Informatics, The Norwegian Radium Hospital, Oslo University Hospital, Oslo, Norway; Dept. of Tumor Biology, Institute for Cancer Research, Oslo University Hospital, Oslo, Norway; Research Support Services Group, University Center for Information Technology, Oslo, Norway; Department of Biosciences, University of Oslo, Oslo, Norway; Department of Cancer Research and Molecular Medicine, Norwegian University of Science and Technology (NTNU), Trondheim, Norway; Centre for Ecological and Evolutionary Synthesis (CEES), Department of Biosciences, University of Oslo, Oslo, Norway

## Abstract

Genome-wide, cell-type-specific profiles are being systematically generated for numerous genomic and epigenomic features. There is, however, no universally applicable analytical methodology for such data. We present GSuite HyperBrowser, the first comprehensive solution for integrative analysis of dataset collections across the genome and epigenome. The GSuite HyperBrowser is an open-source system for streamlined acquisition and customizable statistical analysis of large collections of genome-wide datasets. The system is based on new computational and statistical methodologies that permit comparative and confirmatory analyses across multiple disparate data sources. Expert guidance and reproducibility are facilitated via a Galaxy-based web-interface. The software is available at https://hyperbrowser.uio.no/gsuite

## Introduction

Improvements in sequencing technologies in recent decades have enabled the determination of the DNA sequences of many large genomes as well as their functional interrogation. Genome-wide profiles for a variety of biological features are being systematically generated for a wide range of cell types, often via concentrated efforts by dedicated consortia. The Encyclopedia of DNA Elements (ENCODE) [1] project marked a substantial leap in this respect by making available to the human genomics community a broad collection of cell line-specific data on DNA accessibility and transcription factor binding. The NIH Roadmap Epigenomics Mapping Consortium further contributed a significant amount of additional tissue- and cell-type-specific data to the public domain, including DNA methylation and histone modification profiles for a large number of primary cells that. Kundaje et al. [2] refer to the combined collection of ENCODE and Roadmap data as 127 human reference epigenomes. Most of these datasets are in the form of genomic tracks, i.e. sets of elements anchored to locations in a reference genome, which provide a good foundation for the integration of data representing disparate genomic features.

The widespread utilization of these immense amounts of available data is hampered by a lack of tools providing automatic data integration and sound statistical analysis of large collections of diverse datasets. Frameworks and toolkits such as Bioconductor [3] (R), bedtools [4] (command line), Galaxy [5] and HyperBrowser [6] (web interface) have enabled the robust processing and analysis of genomic tracks with reduced development effort using a variety of interfaces. However, these tools are essentially limited to analyses involving either a single track or a pair of tracks, with no support for the analysis of track collections beyond the trivial concatenation of results per track. For investigations aiming to exploit larger data collections through comparative analyses across epigenomes or across genomic features, no general solutions are available (on any platform). Dedicated solutions do exist for specific applications (e.g., assessing a cell type-specific accessibility of a set of single nucleotide polymorphisms (SNPs) [7, 8] or annotating genomic variants [9, 10, 11, 12]), for specific analytical scenarios (e.g., enrichment analysis of one track against a collection[13]), and for specific basic operations (e.g., calculating the number of base pairs covered by all tracks in a collection [14] or computing the intersection of a collection of tracks with the elements of a single query track [10]). Figure 1 presents these different frameworks and dedicated solutions in context. The lack of comprehensive methodologies leads to ad hoc development of analytical solutions in attempts to answer novel questions that draw on the power of large public or in-house data collections. This may severely limit exploitation of the full potential of current experimental technologies and public data repositories, particularly by research groups with limited bioinformatics resources. Furthermore, the prevalence of ad hoc solutions has a negative impact on reproducibility. A new layer of computational methodology is thus needed to directly approach generic questions formulated in the domain of track collections.

**Figure 1:**
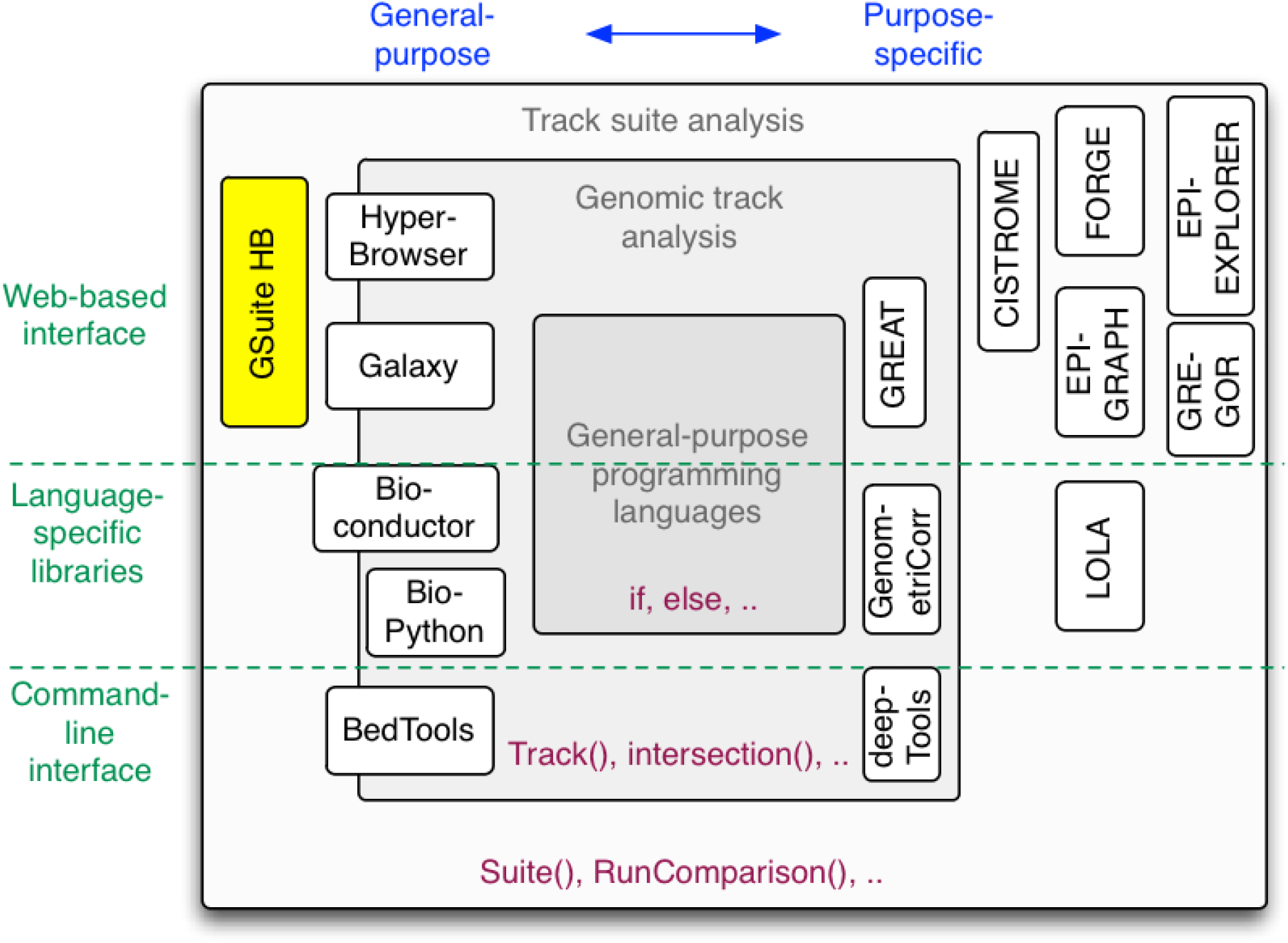
The GSuite HyperBrowser in the context of existing tools and frameworks for genomic track analysis. The codebases of frameworks such as bedtools [4], BioPython [28], Bioconductor [3], Galaxy [5] and the Genomic HyperBrowser [15] add a domain-specific layer on top of general programming languages, providing generic constructs for representing genomic track data and core operations on tracks (including some minimal support for analyzing multiple tracks). The GSuite HyperBrowser codebase is the first general platform to add a new layer of constructs for directly representing collections of tracks and providing core operations (analyses) to be applied to such track collections. Although the functionality of this codebase is provided through a web interface, the codebase is open source, and the same constructs may be used by any other relevant Python-based platform. Also, the underlying approach is general and could be correspondingly implemented in other programming languages. In addition to such general purpose framework, there are a variety of purpose-specific tools for track data. GenometriCorr [29], deepTools2 [30] and GREAT [31] are examples of tools that operate on single/pairs of tracks and support specific analyses or domains. Furthermore, several tools implicitly make use of collections of genomic tracks for analyses in specific domains (e.g., FORGE [8], GREGOR [7] and CISTROME [18]) or for specific types of analyses (e.g., EpiGraph [32], MULTOVL [14], EpiExplorer [33] and LOLA [13].

Here, we present GSuite HyperBrowser, the first comprehensive solution for the analysis of track collections across the genome and epigenome. GSuite HyperBrowser is an open-source, web-based system that enables analysis of a broad array of both hypothesis-driven and data-driven questions that may be posed using large collections of genomic tracks. We focus on questions of a comparative nature, where a track is contrasted to (or analyzed in the context of) other tracks. The intended input is one or more carefully assembled collections of tracks, with the tracks of a collection typically varying along a single dimension of interest. The input could be a collection of tracks for the same histone modification across cell types or a collection of tracks representing different histone modifications in the same cell type. The system uses a formalized representation of track collections and includes tools for compiling new collections from local files or public repositories. Analytical questions may relate to which tracks stand out from such a collection, which tracks of a collection are the most similar to a separate (query) track, or how the occurrence or co-occurrence of elements from individual tracks in the collection varies along the genome. Included within the system is guidance on how these generic questions can be meaningfully interpreted with respect to a specific genomic feature.

## Results

### Overview

The present work is concerned with sets of information elements anchored to specific coordinates in a reference genome, which we refer to as genomic tracks (short form: tracks). A genomic track may denote e.g. the genome-wide set of experimentally determined locations of DNA methylation or DNA binding by a transcription factor. Often, an investigation may involve a carefully selected collection of tracks representing either different genomic features for a single cell type or a single feature for multiple cell types. We refer to collections of tracks selected for a particular analytical purpose as suites of tracks (short: suites).

We define a simple and intuitive tabular format, GSuite, to represent suites of tracks. The GSuite format can represent data at a local or remote server, can include metadata, and can be seamlessly exchanged between individual tools in an analysis workflow. To allow efficient compilation of track suites from a variety of public repositories (like ENCODE and Roadmap Epigenomics) and thus enable integration of disparate data sources, we propose that rather than downloading and reorganizing tracks according to a unified structure, a concept akin to database views is preferable; tracks can be browsed and selected in a unified manner but are retrieved from their respective sources only when a user assembles a track suite.

Even for a pair of tracks, many different questions can be asked regarding their relations [15]. In principle, the number of possible relations that can be queried for multiple tracks grows exponentially with the number of tracks involved. Also, the complexity of defining and interpreting analyses involving multiple heterogeneous tracks is very high. A particularly useful type of question is the comparative assessment of tracks in a suite, where the tracks may be contrasted based on their relation to one another, to a particular separate track or to tracks of another suite. We delineate a set of generic questions that are useful across a broad range of investigations, explore their characteristics, and present a statistical methodology for their resolution. Table 1 lists five of the main questions, along with associated descriptive statistics and hypothesis tests (details provided in Additional file 1). The descriptive statistics can be based on different measures of similarity, and the hypothesis tests can be based on different null models.

**Table 1:**
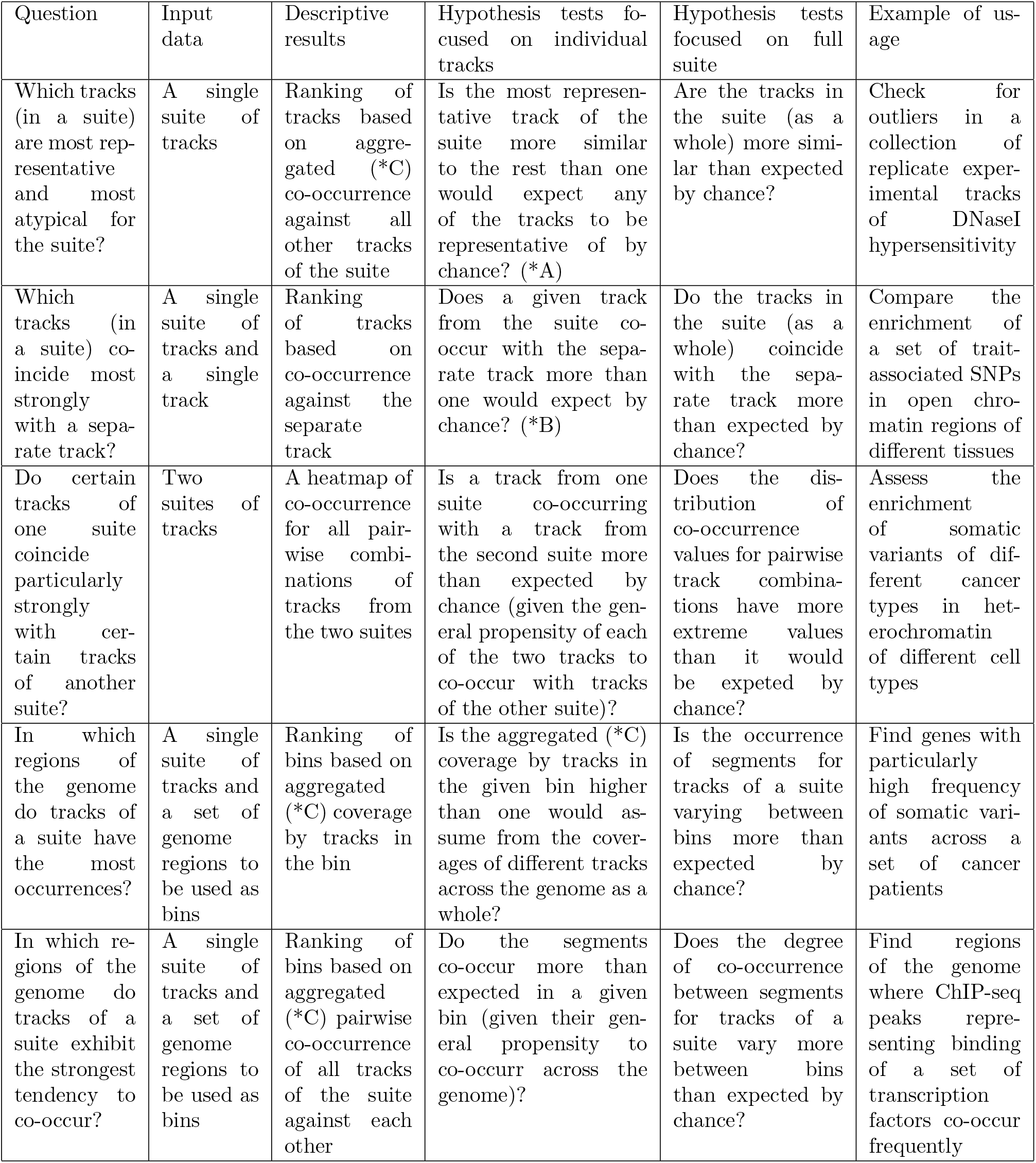
Analytical questions on track collections.

| Question | Input data | Descriptive results | Hypothesis tests focused on individual tracks | Hypothesis tests focused on full suite | Example of usage |
| --- | --- | --- | --- | --- | --- |
| Which tracks (in a suite) are most representative and most atypical for the suite? | A single suite of tracks | Ranking of tracks based on aggregated (*C) co-occurrence against all other tracks of the suite | Is the most representative track of the suite more similar to the rest than one would expect any of the tracks to be representative of by chance? (*A) | Are the tracks in the suite (as a whole) more similar than expected by chance? | Check for outliers in a collection of replicate experimental tracks of DNaseI hypersensitivity |
| Which tracks (in a suite) coincide most strongly with a separate track? | A single suite of tracks and a single track | Ranking of tracks based on co-occurrence against the separate track | Does a given track from the suite co-occur with the separate track more than one would expect by chance? (*B) | Do the tracks in the suite (as a whole) coincide with the separate track more than expected by chance? | Compare the enrichment of a set of trait-associated SNPs in open chromatin regions of different tissues |
| Do certain tracks of one suite coincide particularly strongly with certain tracks of another suite? | Two suites of tracks | A heatmap of co-occurrence for all pairwise combinations of tracks from the two suites | Is a track from one suite co-occurring with a track from the second suite more than expected by chance (given the general propensity of each of the two tracks to co-occur with tracks of the other suite)? | Does the distribution of co-occurrence values for pairwise track combinations have more extreme values than it would be expected by chance? | Assess the enrichment of somatic variants of different cancer types in heterochromatin of different cell types |
| In which regions of the genome do tracks of a suite have the most occurrences? | A single suite of tracks and a set of genome regions to be used as bins | Ranking of bins based on aggregated (*C) coverage by tracks in the bin | Is the aggregated (*C) coverage by tracks in the given bin higher than one would assume from the coverages of different tracks across the genome as a whole? | Is the occurrence of segments for tracks of a suite varying between bins more than expected by chance? | Find genes with particularly high frequency of somatic variants across a set of cancer patients |
| In which regions of the genome do tracks of a suite exhibit the strongest tendency to co-occur? | A single suite of tracks and a set of genome regions to be used as bins | Ranking of bins based on aggregated (*C) pairwise co-occurrence of all tracks of the suite against each other | Do the segments co-occur more than expected in a given bin (given their general propensity to co-occur across the genome)? | Does the degree of co-occurrence between segments for tracks of a suite vary more between bins than expected by chance? | Find regions of the genome where ChIP-seq peaks representing binding of a set of transcription factors co-occur frequently |

The representation, acquisition and analysis of track suites are implemented in a comprehensive, open-source software system, GSuite HyperBrowser. The system builds on the Genomic HyperBrowser [15, 6] and offers a web-based interface powered by Galaxy [5], with several separate tools for the compilation, preprocessing and analysis of track suites (Figure 2). The web interface includes an interactive tutorial to help new users quickly get up to speed with meaningful analyses, guidance for every tool, and a set of thoroughly annotated examples of biological investigations.

**Figure 2:**
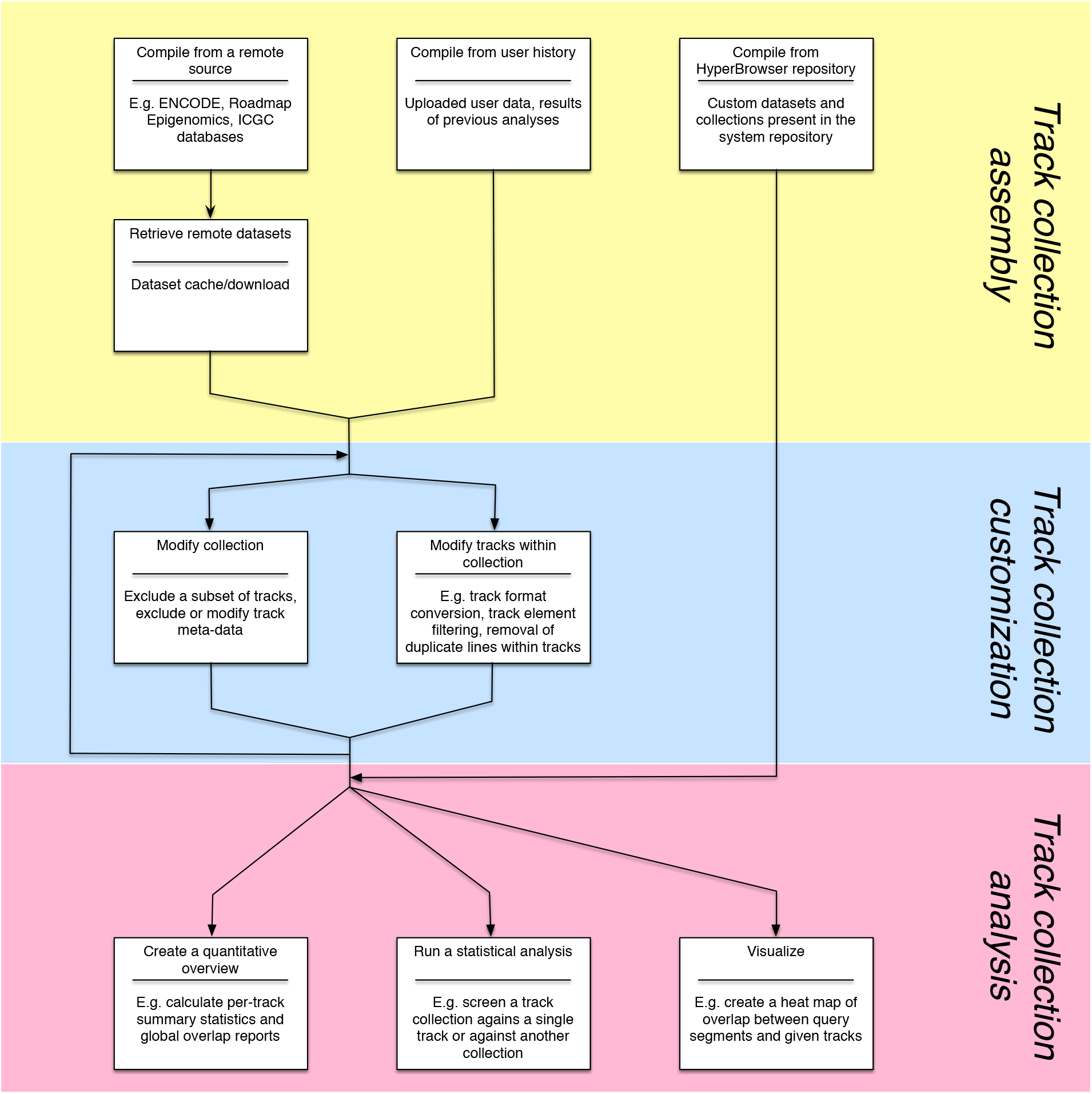
Overview of typical analysis phases and the tools included in the GSuite HyperBrowser system

### Illustrative example

As an illustrative example, consider the exploration of how binding sites for a given transcription factor (TF) co-occur with binding sites of other TFs and with various epigenomic marks. Because TF binding varies between cell types, such an exploration should be conducted in a cell type-specific context. Here, we describe a process for determining the cooccurrence of ChIP-seq peaks for the GATA1 TF versus other TFs and functional epigenomic elements in K562 cells, an established cell line for which abundant experimental data are available. All analysis steps are performed using tools within the GSuite HyperBrowser system. Further details of the analysis and biological interpretations are discussed in Additional file 2.

The first step is to browse available experimental datasets for K562 cells in the ENCODE repository, compile a GSuite file referring all K562 ENCODE tracks and download these to the server (318 tracks). Using tools for GSuite customization, we isolated a single GATA1 track and compiled a suite of the 317 remaining tracks.

We then determined which tracks (in the suite) exhibit the strongest similarity (in terms of peak cooccurrence) with the GATA1 track. The most critical aspect of such an analysis is the precise specification of the measure of similarity (co-occurrence). By selecting the Forbes measure [16], we obtain a ranking of track similarity that is unbiased by the strongly varying number of elements in each track. By performing the analysis in this manner, the transcription factors SMARCA4, SIRT6 and SMARCB1 (Ini1) were identified as high-ranking. These TFs have all been previously reported as relevant for GATA1 (see discussion in Additional file 2).

Because we did not filter out any K562 tracks included in the suite, the ranking includes experimental replicates for GATA1 as well as non-TF datasets such as histone modifications and DNase I accessibility. This provides a broad view of co-occurrence, including indications for TF cooperation, consistency across experimental replicates for the same TF, and the association of GATA1 with different chromatin states. As a confirmatory extension of the analysis, one can examine whether the high-ranked tracks are significantly more similar to GATA1 than the average for all tracks in the suite. This question can be answered by a hypothesis test available in the same tool used to produce the ranking; it uses a test statistic comparing the similarity of each track to the average of the suite. Different null models may be reasonable; for instance, a null model may assume that the data in the whole suite are fixed, whereas the peak locations in the separate track (GATA1) are assumed to be stochastic according to a distribution that preserves the empirical distribution of lengths and distances between the peaks [15]. Because an average across the suite forms part of this test statistic, data for the whole suite are required to compute each single measure, meaning that the analysis is at the integrative multiplicity level (as defined in the section on Classes of multiplicity).

### Representing suites of genomic tracks: the GSuite format

Fundamentally, a collection of datasets is fully defined by a set of references to its constituents. For convenience, a plain text file of Uniform Resource Locators (URLs) for the contained datasets should be valid as a representation of a dataset collection. To further support relevant analyses, the format should permit inclusion of metadata defining important attributes of each individual dataset.

We have defined a simple format that meets these requirements, GSuite. A plain text file of one URL per line is a valid GSuite instance. The format further allows the definition of headers that, among other functions, declare whether the included datasets are available locally or remotely. A tool that downloads datasets referred to by a collection can then iterate through the source GSuite, download each referred file, and replace the URLs with paths to the locally stored files. In addition to the URLs of the tracks, a GSuite file may include tab-separated columns representing metadata values for each dataset. A full definition of the GSuite format is provided in Additional file 3.

### Compiling suites from public repositories

Although repositories such as ENCODE and Roadmap Epigenomics provide free access to large amounts of data, they are not designed for the extraction of large numbers of datasets according to shared characteristics, e.g., extracting large suites of tracks tailored toward a particular analysis. Furthermore, the different repositories do not use a common nomenclature, hindering the integration of related data from several repositories.

A common solution to the integration of data from multiple repositories is to download all data from their respective sources, and construct a meta-repository structured according to a common terminology (e.g., [17, 15, 18]). However, such manual curation and organization is laborious, susceptible to imprecision or misunderstanding, and can easily become outdated. We therefore adopted a different approach to integrate tracks from multiple sources. Rather than downloading and re-organizing genomic tracks, we use a concept akin to database views; users can browse and select remotely located tracks based on metadata, resulting in a list of URLs of the chosen tracks (GSuite). The GSuite can be further modified and shared as a simple text file. The underlying genomic tracks are only downloaded when a user explicitly asks to create a local copy of the data.

As a low-level access point, we provide a single interface for accessing different repositories according to their original (repository-specific) metadata terminology. This interface avoids the loss or misrepresentation of the exact metadata provided by the individual repositories.

We also provide a high-level access point that sacrifices some degree of metadata precision to permit selection of related tracks across sources according to a unified vocabulary (e.g., all tracks for a particular histone modification across repositories). The high-level access point builds on the low-level access point and is based on a curated transformation of individual repository-level vocabularies into the unified vocabulary.

The low-level and high-level access points currently support ENCODE [1], Roadmap Epigenomics [2], the International Cancer Genome Consortium data portal [19] and the NHGRI-EBI GWAS Catalog [20].

### Classes of multiplicity for analyses of track suites

The analysis of multiple tracks ranges from simple repetition of the same computation on each track to analyses in which the tracks are highly intertwined in the computations and interpretations. To better delineate the different levels of integration associated with various analyses, we define the following classes of multiplicity for track suite analyses:

#### Trivial multiplicity

A statistic is computed for each track in a suite, but the computed values are neither compared nor integrated across tracks in the suite of interest. This resulting list of values per track can be convenient for obtaining an overview of a suite. Because it is merely a repetition of computations, it does not introduce any challenges related to multiplicity. An example of trivial multiplicity is to count the number of peaks for each track for transcription factor binding sites in a given cell type.

#### Contrasting multiplicity

A statistic is computed separately for each track of a suite, possibly in relation to reference tracks (outside the suite), with an aim of contrasting (typically ranking) the values computed for each track from the suite. Cooccurrence is typically at the core of the computations. Although the computations are performed separately (as with trivial multiplicity), the aim of comparing the computed values puts additional requirements on the statistics used. As discussed in Additional file 2, measures designed to capture the similarity/co-occurrence of tracks may have biases related to *e.g*., the number of elements in a track. An example of contrasting multiplicity is to evaluate the co-occurrence of binding sites of a selected transcription factor (TF) against each track from a suite of transcription factor ChIP-seq peak tracks ^*^. In this example, using the Forbes measure [16] to assess co-occurrence resulted in a biologically very reasonable ranking of potentially cooperating TFs, whereas the Jaccard measure [21] produced a ranking that appeared severely biased by the number of peaks in each track from the suite.

#### Integrative multiplicity

A statistic is computed based on pair-wise measures across all tracks in a suite. The statistic may be a single value representing the suite as a whole or it may be in the form of one value per track from the suite. For descriptive statistics computed per track, integrative multiplicity implies that the value of a given track will depend on the context of other tracks included in the suite. An example of integrative multiplicity is the computation of how typical each track in a suite is with respect to the suite, i.e., its average co-occurrence with other tracks in the suite. A computational challenge associated with the integrative multiplicity class is that the data for each track are typically used in several parts of the computations. A simple algorithm would thus either need to read the same data repeatedly from physical storage or simultaneously store the data for all tracks in memory. More advanced algorithms based on map-reduce and memoization of intermediate computations would therefore generally be preferable (and are applied in GSuite HyperBrowser).

#### Higher-order multiplicity

A statistic is defined based on higher-order relations (beyond pairwise) between the tracks in a suite, implying that a computation must work on elements from many/all tracks from a particular genomic region simultaneously (a statistic that cannot be subdivided into multiple pairwise across-track computations). An example is the computation of how many base pairs across the genome are associated with open chromatin in more than half of a set of considered cell types (covered by more than half of the genomic tracks of a suite).

### Hypothesis testing

A hypothesis test for multiple tracks investigates whether the aspect of interest for the track or tracks in question is present in the data more/less than what is expected by chance. For all questions in Table 1, we have defined an associated statistical test that can facilitate the assessment of the robustness of the effects observed in the descriptive statistics (Additional file 1).

Statistical tests can be based on parametric distributions or Monte Carlo simulations. Due to the complex structure of a genome, genomic data sets are often not well described by simple parametric distributions. For this reason, simulation has been the preferred choice even for relations involving only a pair of tracks [15, 22]. We have further demonstrated that the simplifying assumptions that are typically required to allow parametric testing on genomic track data will often increase the risk of false-positive findings [23]. Based on such considerations, we find that for the questions of Table 1, the limitations and simplifying assumptions required for parametric testing make Monte Carlo-based simulation a more promising direction.

The following are the main elements of a Monte Carlo-based statistical test: 1) *a test statistic:* a measure that describes the aspect of interest; 2) *a null model:* a model that tracks would follow if generated by chance; 3) *a null distribution:* the distribution of the test statistic when data follow the null model; and 4) *a p-value:* the proportion of the null distribution that is more extreme than the value of the test statistic on the observed (real) data. For statistical testing to be meaningful, a test statistic must be specified that precisely matches a particular aspect (question) of interest and assumes a realistic (relevant) null model.

Our approach follows [15]; we argue that good models can be obtained by preserving some structure from the tracks and by randomizing others. After specifying what we consider relevant null model assumptions, we derive algorithms for sample tracks from a particular null model and compute the test statistic for each simulated track. We observe that the relevant null models (and thus the associated simulation algorithms) are mostly shared between questions and can be divided into the following three categories (described in terms of simulation algorithms):

- Sampling algorithms that treat each track separately. Any sampling algorithm for single tracks can be extended in this manner to suites, e.g., those presented in [15].
- Sampling algorithms that sample elements across tracks from a suite. Track segments (pairs of reference genome coordinates) can be placed in a single pool shared across tracks and sample segments for each track with or without replacements from this pool and with or without preserving the variation of frequency and length of segments across the tracks. A particular challenge with this sampling approach is how to handle intra-track overlap of segments without introducing sampling biases. Further details on alternative sampling algorithms are provided in Additional file 1.
- Sampling algorithms sampling across suites. These fall into the following two types: one type that pools track elements across both tracks and suites and thus represents a (slight) further complication of the previous category and a second type that permutes entire tracks between suites. Further details are provided in Additional file 1.

There is a crucial difference in the interpretation between hypothesis tests at contrasting and integrative multiplicity levels. A statistical test that uses a pairwise track similarity measure as a test statistic and a sampling algorithm that treats each track separately will result in p-values at the contrasting multiplicity level (p-values relate to the null hypothesis for each track from a suite in isolation). Such p-values do not provide information about how a particular track is differentiated from other tracks in a collection, but the p-values of different tracks can be compared to assess the relative confidence. By contrast, if either the test statistic is defined across tracks from the suite or if the sampling algorithm draws elements across tracks, the resulting p-values will be at the integrative multiplicity level. Such p-values may represent null hypotheses related to whole suites or how a given track is differentiated from the remaining tracks in the suite.

### The basic mode as an interactive tutorial of the system

To accommodate a broad range of usage scenarios, the main tools in the GSuite HyperBrowser are defined in a generic and highly customizable manner. Generality of tools and a rich palette of parameter options are often indispensable for appropriate handling of data during the course of an actual project (and often have important consequences for the interpretation of results), but might mean unnecessary complexity for new users who wish to first familiarize themselves with the system. The system therefore includes a dedicated tutorial version of the tool interface, which simplifies the definitions of basic analyses and streamlines the learning experience. This “basic mode” of the system offers a simplified view of a tool’s parameter list, hiding options that are typically sufficiently represented by the default values during initial exploratory test runs by users. Perhaps most importantly, the entry point of the basic mode is a set of interactive analysis examples that illustrate the typical usage of the GSuite tools within particular domains (e.g., the study of genome variation or the study of transcription factor binding). Each example includes detailed instructions for performing a simple integrative analysis and provides relevant datasets necessary for its execution. The examples also offer information regarding generalization of the presented analyses and guidance for utilizing one’s own datasets. Entering and leaving the tutorial mode is possible at any time, which will respectively hide or reveal the full set of parameters defined for each tool.

### Examples of biological investigations using the system

While the interactive tutorial illustrates core analytical approaches for a breadth of biological questions, a full investigation will usually involve its own specific steps for data preparation and supporting analysis. To provide an impression of the variety of aspects that may be involved, we include a set of transparent and reproducible examples of biological investigations using the system. The investigation examples are available under the ‘Examples” tab on the system front page and include an example that reproduces individual findings from the literature (relationship between mutations in a given cancer and cell-specific open chromatin), an example of novel investigations (whether SNPs associated with various diseases are located in miRNA genes), an example of studying experimental biases/artifacts (clustering of tracks associated with different cell types and experimental setups) and an example of studying computational biases (how the exact formula used to measure track similarity has a decisive impact on the results and interpretations).

## Discussion and Conclusions

Reference genomes have allowed a broad range of genomic features to be represented in a uniform manner, which facilitates data integration and the discovery of relations and interplay between various features. With recent initiatives to unravel data from multiple epigenomes (cell-type-specific data for a variety of epigenetic marks), a new layer of computational methodology is needed. Similar to the previous generation of computational tools that allowed a question regarding a genome-scale data set to be resolved through a single operation, the next generation of tools (or an updated version of existing tools) should directly approach questions formulated in the domain of collections of genomic tracks.

The most trivial level of functionality for analyzing data collections, based on iterative, single or pair-wise analysis of genomic tracks, is already available on various platforms for genomic track analysis. More complex solutions regarding track collections have been provided only for specific questions by means of dedicated tools (e.g., LOLA [13]). The analysis of track collections (e.g., analysis across a set of functional elements or cell types) has received little attention in the literature. We present here a first step in this direction.

The present work includes three distinct contributions: 1) a computational and statistical methodology for compiling and analyzing collections of genomic tracks; 2) an implementation of the proposed methodology in the form of a large open-source, integrated software system; and 3) a web-based interface to the developed functionality. The user interface enables meaningful analysis customization by providing expert guidance.

The main approach for the integration of data in the bioinformatics field has been to download data from multiple sources and restructure it according to a uniform hierarchy ([17, 24]). Here, we adopted a different approach by developing solutions to allow users to retrieve data from databases when a specific collection of tracks is needed (instead of downloading and re-organizing data in a general manner in advance). This approach has advantages and disadvantages. Downloading and integrating track collections as needed introduces a delay for users at the time of compilation compared to relying on pre-collected data. This delay is to some degree rectified by a scheme for locally caching data previously downloaded (by any user). The advantage of the chosen approach is that as long as the repositories continue to release their data according to the same protocol, the tool will continuously provide access to all available data in their latest versions. Another strong advantage is the transparency of the approach—users can directly view the URLs at which data were retrieved and the exact time the data were retrieved from a given repository. The currently supported repositories all contain data for the human genome, but the methodology can be readily applied to data connected to any reference genome.

Due to the size and heterogeneity of the genomes of higher organisms, even analyses of single genomic tracks can be complex. Integrative analyses across multiple tracks (typically across cell types or features) add a further layer of complexity. To cope with this complexity, highly customizable tools and extensive user guidance are essential. By developing an integrated software system with a set of robust components for data handling and statistical analysis at the core, we have enabled a range of sophisticated analyses to be performed with limited effort. The developed methodology is accessible to a broad user base via the system’s web interface, which provides inbuilt tool guidance and offers an interactive tutorial with a rich list of domain-specific analysis suggestions. Transparency and reproducibility of analyses are ensured by integration with the Galaxy framework, where data, tool and parameter choices are automatically tracked in the background and any step in the analysis can be repeated with the option of changing the original data or parameters.

The methodology presented here does not cover the full spectrum of analyses that can be envisioned for collections of genomic tracks. First, the current statistics and null models only relate to pure location data (Point and Segment tracks [25]). Extending the work to handle *Valued Points* and *Segments* (e.g., genes with expression values and tracks from case vs. control elements) as well as *Function* tracks (e.g., signal tracks with ChIP-seq intensities) would clearly broaden the range of supported biological investigations. Second, the present methodology is primarily focused on questions that can be reduced to pairwise track relations. Analysis of higher-order relations between functional elements is a very interesting challenge but requires methodological development beyond what is described here. Third, even for the class of analyses considered here, there are many further questions for which statistical methodology would be useful. Fourth, although data from any source can be uploaded to the system, a consistent terminology for track metadata would enable better unified access to track data sources and their content. We believe that the development of a widely accepted ontology for describing biological and experimental characteristics of tracks should be given high priority to ease data integration and avoid misinterpretation of results achieved when employing public data for research. Fifth, experimental data at the single-cell level is rapidly becoming a powerful tool in biomedical research [26, 27]. Although the methodology presented here can be used directly on singlecell data, these data may give rise to a range of additional questions beyond what is considered in the present work. Through a principled methodological approach and implementation based on generic core components, the open-source GSuite HyperBrowser system is prepared for future extensions in a variety of dimensions.

In conclusion, we believe the GSuite HyperBrowser would permit robust and reproducible solutions to a breadth of cases for which ad hoc development is the only current possibility.

## Methods

### System implementation

The GSuite HyperBrowser is an integrated software system written mainly in Python, with extensive use of the NumPy library for efficient data handling, as well as some supporting code in R and Javascript (in total, 170,000 lines of code). The GSuite HyperBrowser makes use of code components from the Genomic HyperBrowser [15] to represent individual tracks and to analyze single tracks and pairwise relations between tracks. The user interface is based on the Galaxy system [5], which ensures robust user and dataset management, and includes features supporting reproducible research. To provide users with a more dynamic user interface, the tools in GSuite HyperBrowser is based upon Galaxy ProTo (https://github.com/elixir-no-nels/proto), an alternative tool definition API for the Galaxy framework. To ensure computational efficiency, track data are preprocessed into an indexed, binary format based upon arrays written consecutively to disk [25], while analysis computations are based on a map-reduce scheme that limits memory requirements and a scheme for memoizing intermediate computations [15].

### GSuite representation

Collections of tracks are represented as lists of references (URLs) with corresponding metadata in the GSuite tabular text format. The system includes robust functionality for composing, modifying and validating collections in this format. The system also includes functionality for crawling and for searching and retrieving data from public repositories. The crawling functionality works similarly to a web crawler, accessing metadata from supported repositories to generate a database of the available datasets in the form of URLs along with metadata accompanying each dataset. This database can then be queried on metadata contents, resulting in a novel GSuite file containing Uniform Resource Identifiers (URIs) to original, remotely stored datasets. Before analysis, remote datasets of a GSuite file can be retrieved and stored locally on the web server in hidden Galaxy history elements, resulting in a transformed GSuite file with custom Galaxy URIs that point to such storage. A caching scheme is also implemented, making sure that the datasets for each unique URI that refers to stable content is only retrieved once. The caching simply stores the Galaxy URI for the first retrieval in a register and makes sure that consecutive retrievals result in the same URI.

### Descriptive statistics and null models

The test statistic needs to be custom-tailored to a particular question. It will thus vary between different questions involving suites of tracks, and will also vary according to slight variations of each question. Still, we find that test statistics for the whole range of questions we have studied can be defined based on a shared hierarchy:

- Pairwise track statistic (T): computes a measure of co-occurrence between a pair of tracks, e.g. the Forbes measure (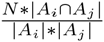, where *A_i_* and *A_j_* are the set of genome locations (bps) covered by two tracks *i* and *j*, while *N* is the size of the genome) [16]. This can be a final per-track result in itself (at the contrasting multiplicity level) or part of a higher order computation.
- Integrative statistic (Q): combines values of T for multiple track pairs. This operates on a structure of track pairs (and corresponding T values), e.g. a single track paired with each other track of a suite. The combination of T values can e.g. be the average, max or min of values of T (e.g., 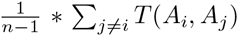), where *n* is the number of tracks in the suite). Analyses based on a Q-statistic are by definition at the integrative or higher-order multiplicity levels.
- Suite statistic (R): Statistic that describes an entire suite. It may combine multiple values of Q. Each Q-value will typically represent a one-to-many computation between tracks in a suite, with the R-value typically representing a many-to-many combination of tracks in a suite. The combination of Q values can e.g. be the average, max or min of values of Q (e.g., 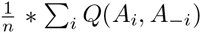). Analyses based on a R-statistic are by definition at the integrative multiplicity level.
- Pairwise suite statistic (S): Statistic that describe the relationship between two suites. Also this statistic may combine multiple values of Q in the same manner as the R-statistic. Analyses based on an S-statistic are by definition at the integrative or higher-order multiplicity level.

Most hypothesis tests in the system are based on Monte Carlo evaluation of p-values, where a particular simulation algorithm produces explicit tracks for the null model and a particular test statistic is used to generate values for the null distribution. Several alternative simulation algorithms are proposed, preserving distinct properties within the scope of individual tracks or across the collection.

Detailed formulas for descriptive and test statistics, as well as detailed sampling algorithms for Monte Carlo evaluation of statistical significance, are provided in Additional file 1.

## List of abbreviations

ENCODE: The Encyclopedia of DNA Elements;
SNP: Single Nucleotide Polymorphism;
TF: Transcription Factor;
URI: Uniform Resource Identifier;
URL: Uniform Resource Locator

## Availability of data and materials

All data and analyses referred to in the manuscript are available from the “Examples” tab on the front page of the GSuite HyperBrowser web page: https://hyperbrowser.uio.no/gsuite

The analyses are available as Galaxy histories, which can be viewed or ‘imported” for further inspection. Full analysis specifications are available through the “run this job again” button present on history elements (this functionality also allows the analyses to be re-run in original or modified form). Data and results can be directly viewed or downloaded.

### Additional files

Additional file 1: A text document describing statistical measures and hypothesis tests for suites of genomic tracks. The document contains detailed formulas and algorithms for statistical methodology used by the GSuite HyperBrowser system. (PDF format, 13 806 KB)

Additional file 2: A text document providing a critical evaluation (on simulated and real data) of how particular choices of similarity measures may influence genome-level analysis results. (PDF format, 1233 KB)

Additional file 3: A text document with a detailed specification of the GSuite file format. (PDF format, 65 KB)

## Footnotes

* As in “Exploring transcription factor co-occurrence using two alternative measures of similarity”, one of the complex example analyses on the GSuite HyperBrower website.

